# NicheScribe: consistent and generalizable annotation of stromal cells in the murine bone marrow niche

**DOI:** 10.64898/2026.09.23.753907

**Authors:** James W. Swann, Jun Hou Fung, Ruiyuan Zhang, Raul Rabadan, Emmanuelle Passegué

## Abstract

Hematopoietic stem and progenitor cells (HSPC) depend on interactions with nearby stromal cells to support their homeostatic functions and mount responses to perturbations. The murine bone marrow (BM) niche can be profiled using single-cell RNA sequencing (scRNA-seq), but there is considerable inconsistency in cell type identification, and uncertainty over how populations can be isolated prospectively by flow cytometry. Here, we produced new scRNA-seq reference datasets using purified populations of murine BM stromal cells, which we used to create a bespoke classifier for stromal cells. Our NicheScribe approach is consistent, accurate, and generalizable to query datasets irrespective of methods used for cell isolation. Moreover, consistent annotation permits comparative analyses of the same stromal cells across different bone types and perturbations like inflammation, neoplasia, and aging. Our work provides new tools for consistent cell type annotation in the murine BM niche and new insights for combined molecular and functional analyses.

**HIGHLIGHTS:**

- New NicheScribe approach for annotation of stromal cells in scRNA-seq datasets
- NicheScribe links molecular state to prospective isolation by flow cytometry
- Different bones have distinct complements of stromal cells
- LepR^+^ mesenchymal stromal cells adopt diverse transcriptional programs upon stimulation

**eTOC BLURB:** Swann et al. develop a computational approach for consistent annotation of stromal cells in murine bone marrow niche, which is applicable to single-cell and spatial transcriptomics. They find that stromal cell frequencies vary in different bones, and that leptin receptor-expressing mesenchymal stromal cells exhibit diverse transcriptional responses to different perturbations.

## INTRODUCTION

Stromal cells in the bone marrow (BM) niche are essential to maintain bone integrity and support the functions of hematopoietic stem and progenitor cells (HSPC), both at steady state and upon aging or a range of perturbations^1–3^. Spatial organization, major cell types, and key receptor-ligand interactions show remarkable conservation between humans and mice^4^, highlighting the importance of the murine BM niche as a key model for understanding regulation of human hematopoiesis. The murine BM niche is increasingly studied using a combination of flow cytometry and single-cell RNA sequencing (scRNA-seq) approaches, which are ideally suited to quantify and profile diverse cell populations isolated in the same cell preparation^5–7^. However, previous studies have generated several different naming schemes for the same stromal cell types, which hampers efforts to perform comparative analyses across conditions. Moreover, it is frequently unclear how molecular states defined in scRNA-seq correspond to cell types that can be isolated by flow cytometry. To address this, we generated a new reference dataset of highly purified murine BM stromal cells according to a validated FACS gating scheme incorporating central marrow (CM) and endosteal (Endo) cell preparations^3^ and used these results to produce a classifier for scRNA-seq data that permits rapid, consistent, and accurate annotation of stromal cell types. We show how this NicheScribe approach can be used to perform comparative analysis of the same cell types across different bones and perturbations, identify shared transcriptional response pathways in the most abundant leptin receptor (LepR)-expressing mesenchymal stromal cell (MSC-L) niche population, and establish an inflammatory MSC-L (iMSC-L) score characteristic of aging and inflammation.

## RESULTS

### NicheScribe consistently annotates BM stromal cells

To provide consistent annotation of stromal cells from the murine BM niche in scRNA-seq data, we designed a classifier with hematopoietic filtering (R1) and 2 layers of stromal cell annotation (R2) (**Figure 1A**). Since stromal preparations are frequently contaminated with CD45^dim^ hematopoietic cells^8^, we first used an extensive atlas of scRNA-seq datasets from whole BM, HSPCs, and both CM and Endo stromal cells in reference R1 to distinguish hematopoietic from non-hematopoietic cells^3,9–11^ (**Figure 1B**), opting to use *Seurat* for integration and label transfer after benchmarking several available software packages (**Figure S1A**). Hematopoietic cells could be annotated effectively using our published *HemaScribe* R package^9^ (**Figure S1B**) and were filtered out before using 2 parallel approaches for annotation of stromal cells. First, we generated a reference R2.1 corresponding to flow cytometric definitions of different stromal cell types by barcoding 10 distinct FACS-purified CM and Endo stromal cell populations using 10X Genomics on-chip multiplexing (OCM) before sequencing (**Figure S1C and S2A**). We used this dataset for annotation of non-hematopoietic cells using integration and label transfer (**Figure 1C**), finding excellent correspondence between predicted R2.1 cell labels and ground truth identities encoded in OCM barcodes (**Figure S2B,C**), as well as expected distributions of cells according to the origin of the cell preparation in samples of CM and Endo origin (**Figure S2D**). Moreover, the R2.1 classification captured stromal cells at similar frequencies in scRNA-seq data to those observed by flow cytometry in matched samples (**Figure 1D**), showing successful alignment of these methodologies. In parallel, we also produced a reference R2.2 corresponding to transcriptomic annotation generated by curation of key marker genes for different stroma cell types using published atlases of mouse BM mesenchymal^3^ and endothelial^12,13^ cells (**Figure S3A-C**). Importantly, this R2.2 reference incorporates molecular heterogeneity of several cell types that cannot currently be appreciated by flow cytometry^2,3^ (e.g., committed fibroblast progenitors [FPr] within the FACS-defined population of Sca-1-expressing MSCs [MSC-S]) (**Figure S2C**), or their CM vs. Endo location (e.g., osteoblast-biased [Osteo] MSC-L and type H endothelial cells [hEC1] found in both locations) (**Figure S2D and Figure S3C**). Collectively, this novel NicheScribe approach provides multiple annotation layers that separates stromal from hematopoietic cells and allows accurate identification of mesenchymal and endothelial BM niche populations (**Table 1**). This method is optimized for matched analyses between molecular and FACS-purified populations while also incorporating recent advances in the field, and all functions are freely available in the R package *NicheScribe* (**Resource Availability**).

**Figure 1.**
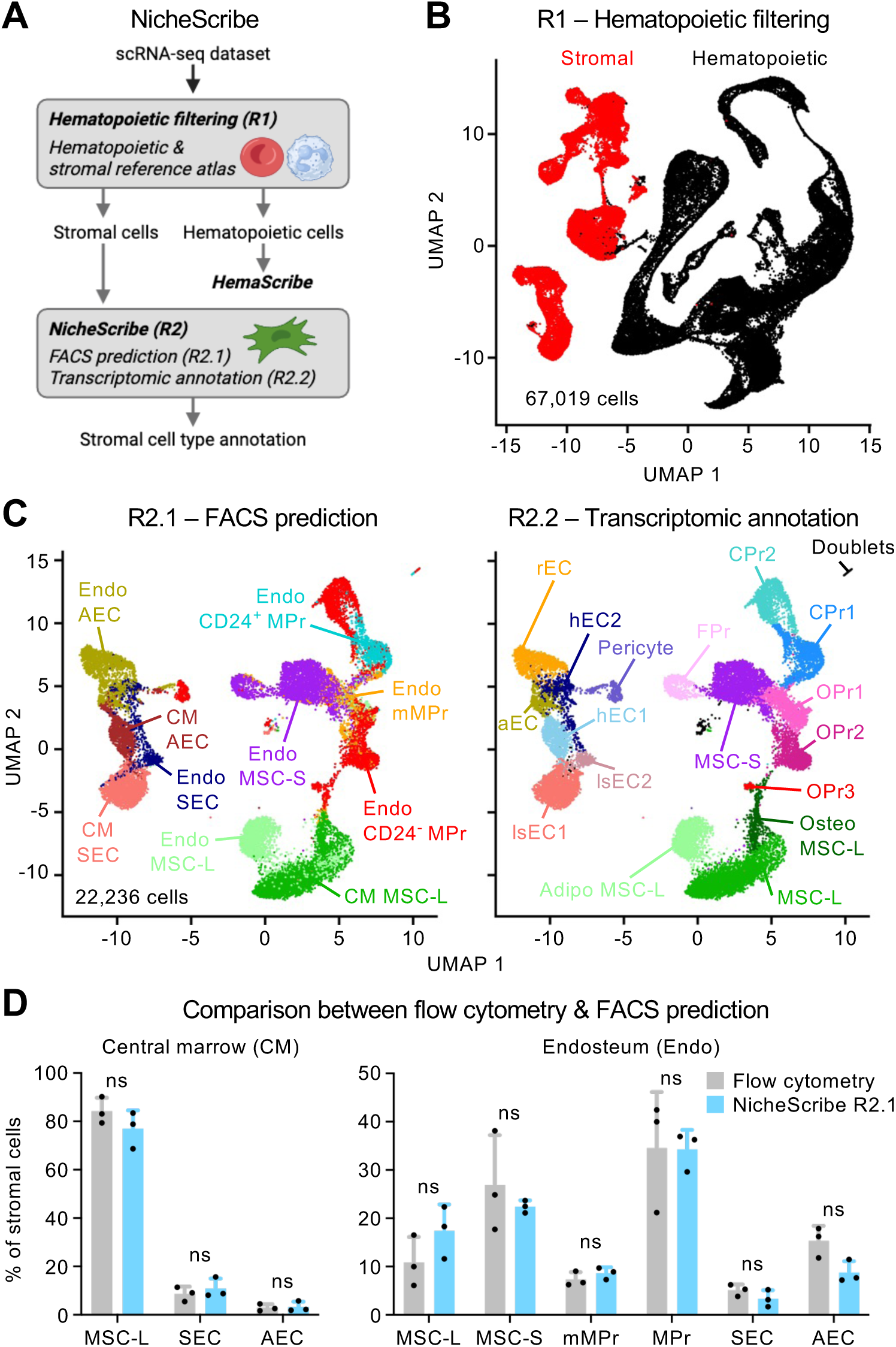
NicheScribe approach for BM stromal cell annotation. (A) Strategy for multilayer annotation of murine BM stromal cells in 10X Genomics scRNA-seq data, beginning with reference (R)1 for filtering hematopoietic cells, followed with R2 for stroma cell type annotation based first on on-chip multiplexing (OCM) of purified populations (R2.1) and then on transcriptomic annotation (R2.2). (B) Uniform manifold and approximation projection (UMAP) of R1 reference data colored for stromal and hematopoietic cells. (C) UMAPs showing R2.1 (left) and R2.2 (right) reference datasets, colored by stromal cell type. AEC or aEC, arterial endothelial cells; CM, central marrow; CPr, chondroblast progenitor; Endo, endosteal; FPr, fibroblast progenitor; hEC, type H endothelial cell; lsEC, type L sinusoidal endothelial cell; (m)MPr, (multipotent) mesenchymal progenitor; MSC-L, leptin receptor-expressing mesenchymal stromal cell with Adipo MSC-L, adipocyte-biased MSC-L, and Osteo MSC-L, osteoblast-biased MSC-L; MSC-S, Sca-1-expressing mesenchymal stromal cell; OPr, osteoblast progenitor; rEC, type R endothelial cell; SEC, sinusoidal endothelial cell. (D) Bar plots showing frequency of indicated cell types in the same samples analyzed by scRNA-seq data using R2.1 transcriptomic annotation and flow cytometry. Points represent biological replicates obtained from pooled cells isolated from 2-4 mice. Bars show mean ± S.D.; *P-values* from two-way ANOVA with Sidak’s *post hoc* test; ns, not significant. See also Figures S1, S2, and S3 and Table S1.

**Table 1.** Correspondence of stromal cell annotations across NicheScribe layers and studies. Harmonization of mouse BM stroma population naming systems across datasets. Type H and Type L ECs showed 2 distinct clusters in NicheScribe transcriptomic annotation (R2.2), which largely aligned with CM and Endo origin, hence the different numbers. Note that other cell types could be present in the bone and/or BM niche but are not captured in NicheScribe annotation owing to loss during cell preparation (e.g., adipocytes, neurons) or failure to release from bone during digestion (e.g. osteoblasts, osteocytes). Indicated most similar human cell types are derived from cell types with highest levels of correlation in gene expression. See also Figures S2, S3, and S5.

### NicheScribe provides consistent and generalizable stromal annotations

To demonstrate the usefulness of our new classifiers, we first re-annotated a published stromal dataset labelled by assigning names to cell types discovered by unsupervised clustering^6^. We found concordance between NicheScribe and original annotations for many populations, but we also found that clusters named ‘Fibroblast 1-5’ actually contained populations committed to different lineages, including true fibroblasts (‘Fibroblast 2’), chondroblasts (‘Fibroblast 4’), and upstream MSC-S (‘Fibroblast 1’) (**Figure 2A**). We then applied NicheScribe to scRNA-seq datasets generated from Endo preparations of different mouse bones (femur, humerus, pelvis, and tibia) to address bone heterogeneity (**Figure 2B**) and the emerging idea that different bones could harbor distinct BM niches^14,15^. This revealed a strikingly different stromal cell composition in the tibia, with a marked bias towards MSC-S and FPr among mesenchymal populations (**Figure 2C**), which was also apparent by flow cytometry (**Figure S4A**), as well as enrichment for type H vessels among endothelial cells (**Figure S4B**). In contrast, the femur and humerus showed a greater tendency towards chondroblast progenitors (CPr1-2), arterial endothelial cells (AEC), and type R endothelial cells (rEC), with the pelvis displaying some intermediate features more similar to the tibia in the mesenchymal compartment and the femur/humerus in the endothelial compartment. Analysis of predicted fate in upstream MSC-S from each bone using *FateID*^16^ revealed corresponding biases towards FPr fate in the tibia and pelvis, and greater bias towards OPr/CPr fate in the femur and humerus (**Figure 2D**). Interestingly, a previous study found more vigorous response to granulocyte colony stimulating factor (G-CSF) in the tibia than in either sternum or humerus^15^, suggesting that the specific cell composition of the tibia could create a BM microenvironment more conducive to emergency myelopoiesis. To determine how NicheScribe might provide insight on spatial localization of BM stromal cells, we analyzed a published spatial transcriptomics dataset from the murine femur generated using 10X Visium HD technology^17^. In these data, we observed the expected localization of Endo osteoblast progenitors (OPr) and Osteo MSC-L along the endosteal surfaces of the diaphysis, as well as in the trabecular bone of the epiphysis (**Figure 2E**). Additionally, Endo MSC-S, FPr, and CPr were all located in adjoining layers at the joint surface. Since Visium spots overlap multiple cells in the dense CM region, we used the R2.2 reference to perform cell type deconvolution using *CARD*^18^, confirming preferential localization of Endo MSC-S at the joint surface, and MSC-L and type L sinusoidal endothelial cells (lsEC1) throughout the marrow in CM/Endo mixed (Mix) locations (**Figure S4C**). Finally, we used the NicheScribe reference data to evaluate the similarity of molecular profiles between human^4^ and murine stromal cell types, finding that gene expression of murine MSC-L was highly correlated with that of human LepR^+^ subsets of APOD+, THY1+, Adipo-, and Osteo-MSC, whereas murine MSC-S and downstream Endo mesenchymal cells were more similar to human Fibro-MSC (**Figure S5 and Table 1**). Together, these results show that NicheScribe is broadly applicable to stromal datasets, irrespective of the originating laboratory and bone type, can be combined effectively with spatial information, and can illuminate conserved features of stromal niche composition across species.

**Figure 2.**
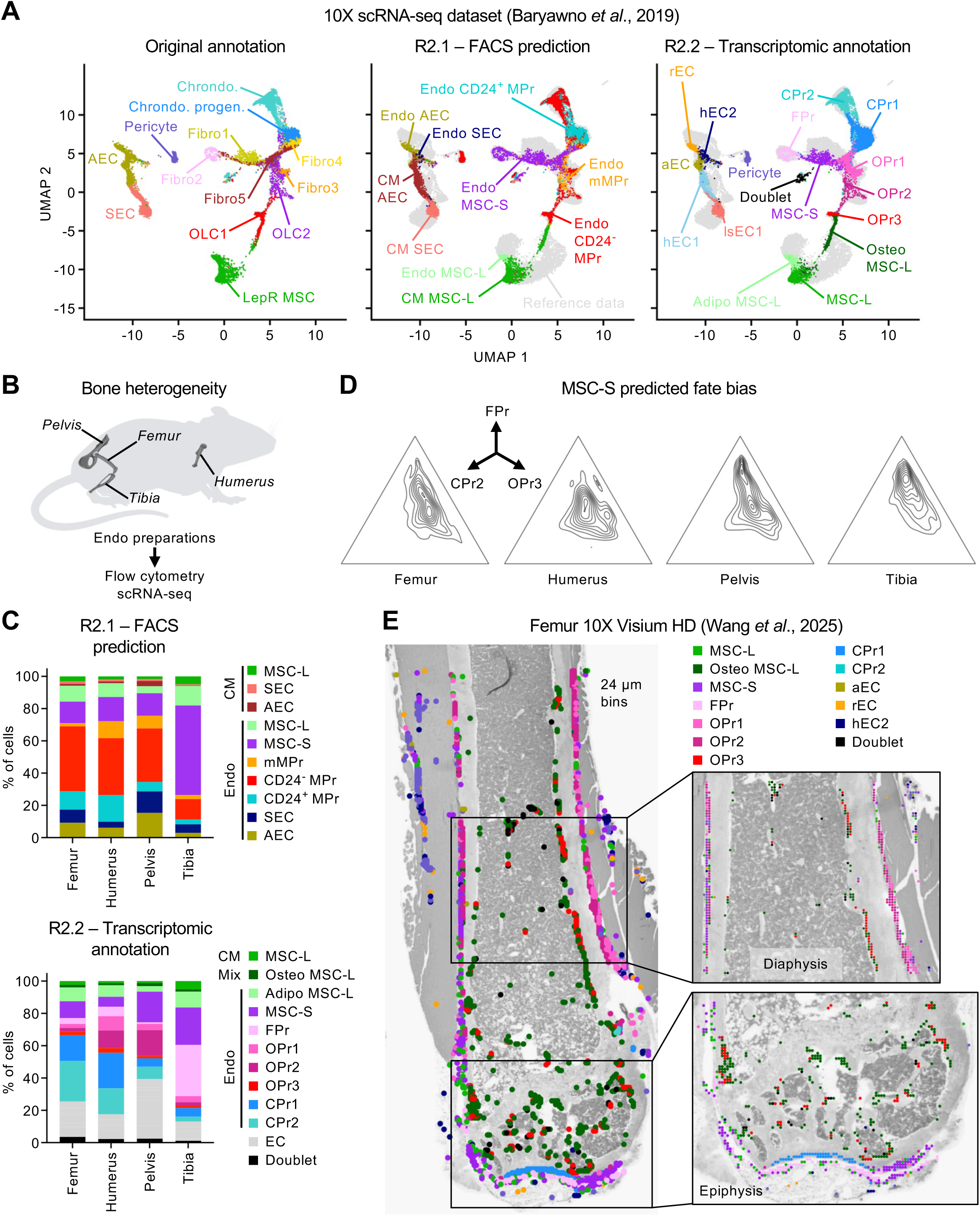
NicheScribe is generalizable to data generation methods and technologies. (A) UMAPs showing original annotation (left) and NicheScribe R2.1 and R2.2 annotations for published scRNA-seq of BM stromal cells. (B) Scheme of the individual bones used for generation of Endo cell preparations for scRNA-seq analyses via OCM. (C) Bar charts showing frequencies of indicated stroma cell types annotated with NicheScribe R2.1 (top) and R2.2 (bottom) in different bones. (D) Ternary plots showing predicted fate bias for MSC-S from indicated bones, using *FateID*. (E) Visium HD of the murine femur, with spots annotated with NicheScribe R2.2 above grayscale hematoxylin and eosin image. Magnified regions show diaphysis and epiphysis. See also Figure S4 and S5.

### NicheScribe facilitates new comparative analyses of stromal cells

We reasoned that consistent annotation of the same cell types across datasets could catalyze new biological insights. To test this, we analyzed published stromal cell data from mice exposed to the cytotoxic drug 5-fluorouracil (5FU)^7^, the viral mimic polyinosinic:polycytidylic acid (pIC)^3^, myeloproliferative neoplasia (MPN) induced by retroviral overexpression of a mutant form of the thrombopoietin receptor *Mpl*^19,20^, and acute myeloid leukemia (AML)^6^ caused by genetic knock-in of an MLL-AF9 fusion transgene^21^ (**Figure 3A**). Comparison of differential abundance of mesenchymal cells using *MELD*^22^ showed that all perturbations caused depletion of some endosteal populations, though with considerable variation in the specific subgroups affected (**Figure 3B and Figure S6A**). Interestingly, 5FU, pIC, and MPN all caused depletion of the most mature osteoblast progenitors (OPr3), suggesting that these stimuli might impair new bone formation, whereas AML caused accumulation of OPr3 and CPr2, consistent with the known effect of AML blasts to stimulate osteoblastic differentiation from upstream MSCs to create a hospitable endosteal niche^23,24^. While the frequency of the core CM MSC-L population showed little change across perturbations, 5FU decreased both Osteo and adipocyte-biased (Adipo) MSC-L subsets, whereas MPN development expanded the Adipo MSC-L subset. Notably, MSC-L identity in high-dimensional space was affected by all perturbations, even if abundance did not change significantly, suggesting altered activation states in MSC-L (**Figure 3B**). To explore this, we integrated stromal cell data and performed consensus non-negative matrix factorization (cNMF)^25^ in MSC-L (**Figure 3C**), identifying 20 factors named M1-20 (**Figure S6B and Tables S1 and S2**). Further analysis showed that 10 factors varied by perturbation (**Figure 3D**), whereas the remaining factors displayed little variation or represented technical artefacts (**Figure S6C**). Among the factors that varied by perturbation, we identified a molecular process M1 of normal MSC-L activity marked by genes encoding bone sialoprotein (*Ibsp*) and trophic factors like angiopoietin-like protein 1 (*Angptl1*), which was suppressed by all perturbations (**Figure 3E**). Interestingly, MPN, and to a lesser extent 5FU, induced a shared molecular process M11 representing myofibroblast differentiation (**Figure 3F**), which resembled a transcriptional program already observed in another murine MPN induced by thrombopoietin overexpression^26^, and which was characterized by genes like α2 smooth muscle actin (*Acta2*), transgelin (*Tgln*), and the secreted factor galectin-1 (*Lgals1*) that has been implicated as a direct driver of MPN-induced BM fibrosis^19^. Exposure to 5FU but not MPN or any other perturbation induced a molecular process M4 of adipogenesis (**Figure S6D**), which resembles changes observed in MSC-L after irradiation^27^ and which is essential for hematopoietic recovery from ablation^28^. Finally, and in line with our previous results, pIC but no other perturbations induced a molecular process M8 of type I interferon (IFN) transcriptional response (**Figure 3G**)^3^. Collectively, these data show how consistent identification of the same cell types across scRNA-seq datasets can uncover shared and unique patterns of cell frequency change and transcriptional activation.

**Figure 3.**
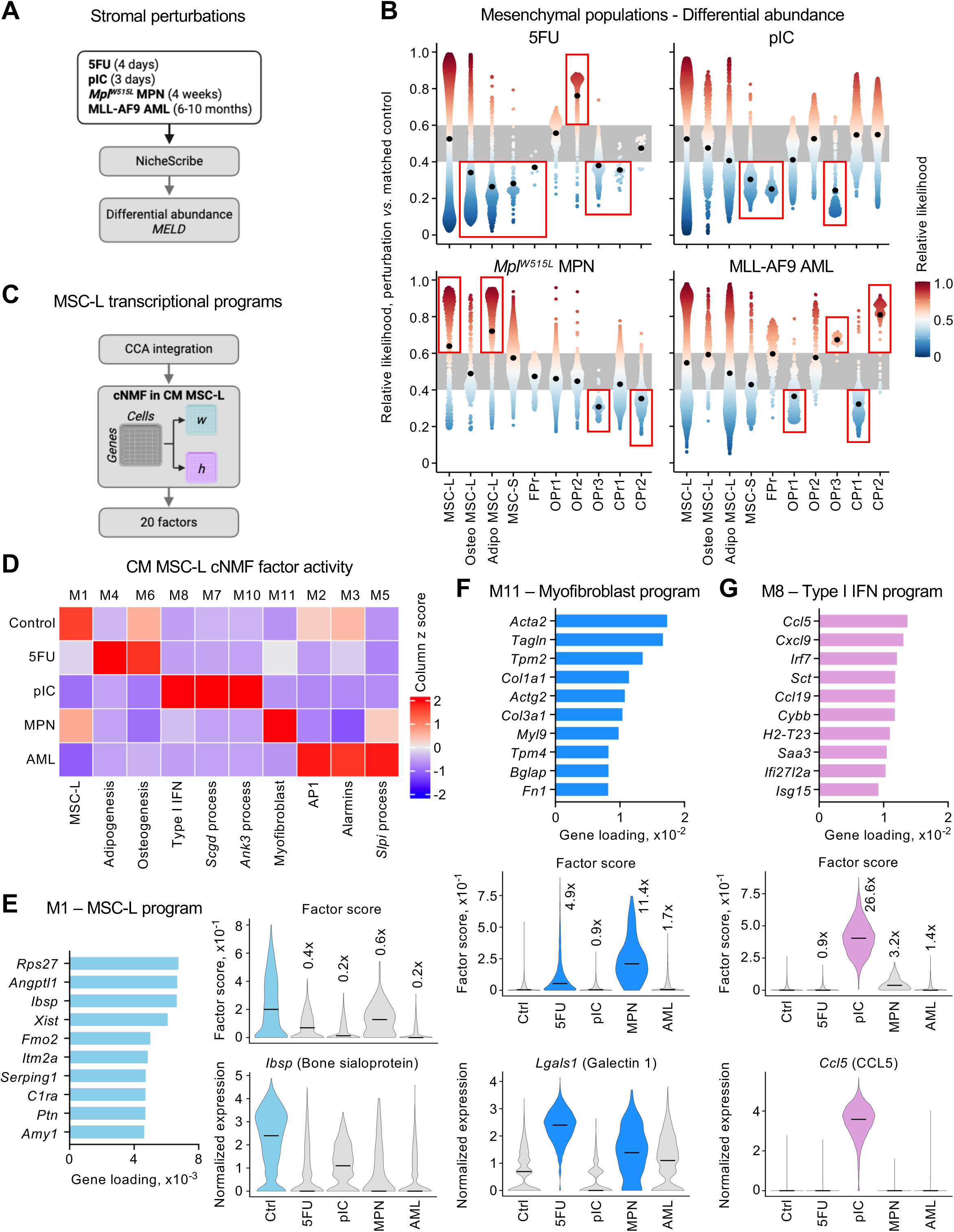
NicheScribe permits new insight into MSC-L transcriptional responses to perturbation. (A) Outline for published perturbation scRNA-seq datasets annotated with NicheScribe and tested for differential abundance with *MELD* using matched control sample for each perturbation. 5FU, 5-fluorouracil at day 4; pIC, polyinosinic:polycytidylic acid at day 3; *Mpl^W515F^* MPN, thrombopoietin receptor mutant myeloproliferative neoplasia at 4 weeks; MLL-AF9 AML, MLL-AF9-induced acute myeloid leukemia at 6-10 months. (B) Beeswarm plots showing relative likelihood of mesenchymal cells annotated with NicheScribe R2.2 in high dimensional neighborhoods. Points represent individual cells and black dots show mean. Values >0.6 indicate increase in relative abundance, <0.4 indicate decreased abundance, and values of 0.4-0.6 (shaded area) indicate unchanged abundance. (C) Outline for investigation of transcriptional response programs in CM MSC-L by canonical correlation analysis (CCA) integration followed by consensus non-negative matrix factorization (cNMF) for 20 factors. (D) Heatmap showing relative activity of 10 selected cNMF factors that varied across perturbations, scaled by column. IFN: interferon. (E-G) Description of selected cNMF factors, with bar charts showing top 10 genes loading for each factor, violin plots showing factor score in individual cells across perturbations annotated with fold change in mean value compared to control, and violin plots for an example gene in each factor. Factors of interest included M1 normal MSC-L program (E), M11 myofibroblast program (F), and M8 type I IFN program (G). Lines in violin plots show median values. See also Figures S6 and S7 and Tables S2, S3, and S4.

### MSC-L inflammation increases with aging

We reported previously that MSC-L can adopt an inflammatory state (iMSC-L) when exposed to type I IFNs, leading to surface expression of Sca-1^[3]^. To provide a quantitative readout of iMSC-L state, we treated mice with pIC for 3 days and then isolated pIC-exposed CM and Endo MSC-L for OCM-based 10X scRNA-seq (**Figure S1C**). Upon comparison of pIC-exposed to steady state MSC-L, we selected the top 100 most significantly upregulated genes and used these to score query datasets using *UCell*^29^ (**Figure S7A,B and Table S3**). We showed previously that aging causes the emergence of the iMSC-L state along with expansion of MSC-L and depletion of most endosteal progenitors^2^, which was confirmed by NicheScribe annotation and quantification of differential abundance of mesenchymal cells using *MELD* (**Figure S7C**). By applying the iMSC-L score across our own^3^ and independently generated aging time courses^30,31^, we confirmed significant upregulation of the inflammatory state in MSC-L over time (**Figure S6D**), highlighting this central feature of BM niche inflammaging. Collectively, this iMSC-L score combined with multilayer annotation in the *NicheScribe* R package provides a comprehensive toolkit for analysis of the murine BM niche.

## DISCUSSION

To improve consistency and accuracy of molecular analyses of the murine BM niche, we developed NicheScribe, which provides comprehensive annotations for different stromal cell populations. These annotations can be linked to flow cytometric definitions to facilitate paired molecular and functional analyses of the same cell types and to allow new insights into the nature of BM niche and its heterogeneity between bones, including the unique cell composition of the tibia that might be more permissive to emergency myelopoiesis^15^. We and others have shown previously that MSC-L are responsive to extrinsic stimuli like cytokines and viral infections^3,32^, and our current analyses demonstrate that MSC-L transcriptional responses are highly diverse and context-specific. Interestingly, a program of myofibroblast differentiation was observed with both MPN disease and cytotoxic 5FU drug exposure, suggesting that malignant cells might co-opt a natural regenerative mechanism to remodel the BM niche, ultimately resulting in hematopoietic failure. Conversely, induction of an adipogenesis program by 5FU and irradiation seems to be essential for hematopoietic recovery from myeloablation. These data suggest that induction of specific transcriptional programs in MSC-L could have dramatic effects on disease phenotypes, and future studies facilitated by our new *NicheScribe* tool kit can now investigate how these fundamental gene expression programs in MSC-L and other cell types are regulated, with the goal of mitigating harmful stromal responses.

## ACKNOWLEDGEMENTS

We thank M. Kissner for management of the CSCI Flow Cytometry Core facilities, and all members of the Passegué and Rabadan laboratories for critical insights and suggestions. J.W.S. was supported by Damon Runyon Cancer Research Foundation DRG-2493-23 (William Raveis Family Fellowship). This work was funded by NIH R01CA184014 and R35HL135763 to E.P., NIH R35CA253126 to R.R., NIH P01CA285250 to E.P. and R.R., and supported in part through the NIH Cancer Center Support Grant P30CA013696 to CUIMC.

## AUTHOR CONTRIBUTIONS

Conceptualization, J.W.S., J.H.F., R.Z., and E.P.; methodology, J.W.S., R.Z., and J.H.F.; investigation, J.W.S., R.Z., and J.H.F.; visualization, J.W.S., R.Z., and J.H.F.; funding acquisition, E.P. and R.R.; project administration, E.P.; supervision, E.P. and R.R.; writing – original draft, J.W.S. and E.P.; writing – review & editing, J.W.S., J.H.F., R.Z., R.R., and E.P.

## STAR METHODS

## Resource Availability

### Lead Contact

Further information and requests for resources and reagents should be directed to and will be fulfilled by the lead contact, Emmanuelle Passegué.

### Materials Availability

This study did not generate new unique reagents.

### Data and code availability

Functions for NicheScribe annotation of cells in scRNA-seq datasets are available in the R package *NicheScribe* with supporting reference data, available for download at <u>github.com/RabadanLab/NicheScribe</u>. The code for *NicheScribe* is archived at Zenodo, 10.5281/zenodo.22922010. Any additional information required to re-analyze the data reported in this paper is available from the lead contact upon request.

## Experimental Model and Study Participant Details

### Animals

All animal experiments were conducted at the Columbia University Irving Medical Center (CUIMC) in accordance with approved Institutional Animal Care and Use Committee (IACUC) and in compliance with all relevant ethical regulations. Wild type (WT) C57BL/6J (CD45.2^+^) mice were purchased from the Jackson Laboratory and bred in house. Mice were 8 to 12 weeks of age when used for experiments, and no specific randomization or blinding protocol was used with respect to the identity of experimental animals. Animal facilities were maintained at 22□±2□°C and 50□±10% relative humidity on a 12–12□hours light–dark cycle, and mice were given *ad libitum* access to Purina LabDiet Rodent Feed and acidified water. Mice were euthanized by CO_2_ asphyxiation followed by cervical dislocation.

## Method Details

### In vivo assays

To harvest inflammatory MSC-L (iMSC-L), 10 mg/kg polyinosinic:polycytidylic acid (pIC, Cytiva) dissolved in sterile PBS was injected intraperitoneally on day 0 and day 2, and mice were euthanized on day 3.

### Central Marrow (CM) and Endosteal (Endo) stromal cell preparation

For each mouse, both femurs, tibiae, hemipelves, and humeri were dissected and thoroughly cleaned. To isolate CM stromal cells, the proximal and distal epiphyses were removed from 1 femur. Intact marrow plugs were flushed with Hank’s balanced saline solution (HBSS) without calcium or magnesium into 5 ml polypropylene tubes by inserting a 3 ml syringe with 22G needle into the distal end of the femur. The plugs were digested with 1 ml of a solution of 3 mg/ml type I collagenase (Worthington) dissolved in HBSS for 10 min at 37°C with 110 rpm shaking. The tube was then vortexed briefly and the supernatant was removed into a new tube through 100 μm mesh. A further 1 ml of 3 mg/ml type I collagenase solution was then added to the marrow plug and the digestion repeated. After the second incubation, the plug was dissociated by pipetting up and down with a P1000 pipette before passing the cell suspension through 100 μm mesh into the tube containing the previous digestion supernatant. To isolate Endo stromal cells, the flushed femur was combined with the other bones, gently crushed up to 5 times using a mortar and pestle, and thoroughly washed with 5 ml HBSS to remove the non-adherent BM cells. The crushing and washing were repeated before the bone chips were collected in a 15 ml polypropylene tube and digested with 3 ml of 3 mg/ml type I collagenase solution for 1 hour at 37°C with 110 rpm shaking. After digestion, the tube was vortexed briefly and the cell suspension was filtered through 100 μm mesh into a new tube. For cell suspensions acquired for both stromal compartments, red blood cells were removed by adding 1 ml ACK lysis buffer (150 mM NH_4_Cl and 10 mM KHCO_3_) and incubated on ice for 3 minutes before washing with HBSS containing 4% fetal bovine serum (HI FBS, Gibco). The cell suspensions were counted using a Vicell automated cell counter (Beckman Coulter). To prepare Endo stromal cells from specific bones, only the 2 bones of indicated types (e.g., femur, humerus, pelvis, tibia) from single mice were crushed and washed for 1 round, then all other processing steps were performed as described above for Endo stromal cell preparation.

### Flow cytometry of stromal cells

For both profiling and sorting of stromal cells from CM and Endo locations, the cell preparations were stained with CD45-APC/Cy7 (BD, 557659; 1:400), Ter119-PE/Cy5 (Invitrogen, 15-5921-83; 1:400), Sca-1-AF700 (eBioscience, 56-5981-82; 1:800), CD31-BV421 (BioLegend, 102423; 1:200), CD105-BV786 (BD, 564746; 1:200), CD51-PE (BD, 551187; 1:100), LepR-biotin (R&D, BAF497; 1:100), CD34-FITC (eBioscience, 11-0341-85; 1:25), CD24-BV510 (BioLegend, 101831; 1:400), CD71-BUV395 (BD, 740223; 1:400) and PDGFRα-PE/Cy7 (Invitrogen, 25-1401-82; 1:100) for 30 minutes on ice. Cells were then washed and stained with Streptavidin-APC (BioLegend, 405207; 1:400) for 15 minutes on ice. Cells were finally washed and resuspended in HBSS with 4% FBS and 1 μg/ml propidium iodide, and then analyzed or sorted on 4-way purity mode on a FACSAria II SORP (CUIMC). Data collection was performed using FACSDiva (v.9) and analysis was performed using FlowJo (v.9/v.10).

### 10X On-Chip Multiplexing (OCM) for stromal cells

Stromal cell populations from different location, bones, or flow cytometry gates were sorted into individual 1.5 ml tubes containing 300 μl of 50% FBS and 50% HBSS, targeting 2,000 cells per population. The following cell populations were isolated (**Figure S1C**):

| Sequence | Cell population | Location | Flow cytometry gate<br>(from Lineage<sup>-</sup>/CD45<sup>-</sup>/CD71<sup>-</sup>) | Cell number<br>collected |
| --- | --- | --- | --- | --- |
| 1 | CM MSC-L | CM | CD31<sup>-</sup>/CD51<sup>+</sup>/Sca-1<sup>-</sup><br>/LepR<sup>+</sup>/PDGFRα<sup>+</sup> | 2000 |
|  | CM SEC | CM | CD31<sup>+</sup>/Sca-1<sup>low</sup>/CD105<sup>high</sup> | 2000 |
|  | CM AEC | CM | CD31<sup>+</sup>/Sca-1<sup>high</sup>/CD105<sup>low</sup> | 2000 |
|  | Endo MSC-S | Endo | CD31<sup>-</sup>/CD51<sup>+</sup>/Sca-1<sup>+</sup> | 2000 |
| 2 | Endo MSC-L | Endo | CD31<sup>-</sup>/CD51<sup>+</sup>/Sca-1<sup>-</sup><br>/LepR<sup>+</sup>/PDGFRα<sup>+</sup> | 1374 |
|  | Endo SEC | Endo | CD31<sup>+</sup>/Sca-1<sup>low</sup>/CD105<sup>high</sup> | 2000 |
|  | Endo AEC | Endo | CD31<sup>+</sup>/Sca-1<sup>high</sup>/CD105<sup>low</sup> | 2000 |
| | Endo mMP<sub>r</sub> | Endo | CD31<sup>-</sup>/CD51<sup>+</sup>/Sca-1<sup>-</sup>/LepR<sup>-</sup><br>/PDGFRα<sup>+</sup> | 1607 |
| 3 | Endo CD24<sup>-</sup><br>MP<sub>r</sub> | Endo | CD31<sup>-</sup>/CD51<sup>+</sup>/Sca-1<sup>-</sup>/LepR<sup>-</sup><br>/PDGFRα<sup>-</sup>/CD24<sup>-</sup> | 2000 |
| | Endo CD24<sup>+</sup><br>MP<sub>r</sub> | Endo | CD31<sup>-</sup>/CD51<sup>+</sup>/Sca-1<sup>-</sup>/LepR<sup>-</sup><br>/PDGFRα<sup>-</sup>/CD24<sup>+</sup> | 2000 |
| | CM iMSC-L | CM | CD31<sup>-</sup>/CD51<sup>+</sup>/PDGFRα<sup>+</sup> | 2000 |
| | Endo iMSC-L | CM | CD31<sup>-</sup>/CD51<sup>+</sup>/Sca-1<sup>low</sup>/LepR<sup>+</sup>/PDGFRα<sup>+</sup> | 1391 |
| 4 | Femur Endo | Endo | Lineage<sup>-</sup>/CD45<sup>-</sup>/CD71<sup>-</sup> | 2000 |
|  | Tibia Endo | Endo | Lineage<sup>-</sup>/CD45<sup>-</sup>/CD71<sup>-</sup> | 2000 |
|  | Pelvis Endo | Endo | Lineage<sup>-</sup>/CD45<sup>-</sup>/CD71<sup>-</sup> | 2000 |
|  | Humerus Endo | Endo | Lineage<sup>-</sup>/CD45<sup>-</sup>/CD71<sup>-</sup> | 2000 |

Following isolation, cells were rested for 1 hour on ice then pelleted at 350 x *g* for 5 minutes at 4°C before the supernatant was removed down to a volume of 10 μl. The samples were loaded onto 10X OCM chips in groups of 4 according to the sequence from the table above. GEM generation and 3’ RNA library preparation were performed according to 10X Genomics protocol CG000768 Rev B, targeting 2000 cell data recovery. RNA libraries were pooled 1:1:1:etc, sequenced on an Illumina NovaSeq X. Library concentrations and fragment sizes were evaluated using Qubit dsDNA HS assay kit (ThermoFisher Scientific) and TapeStation D5000 DNA ScreenTape analysis (Agilent). Fastq files were aligned to the murine mm10 genome using *Cellranger*, v.9.0.

### NicheScribe

We curated multiple scRNA-seq measurements of murine BM to train our *NicheScribe* classifier (LK: GSM7712015, GSM8969792, and GSM9005917; LSK: GSM9005916 and GSM8969793; whole BM: GSM9005914; stromal cells: GSM8505733, GSM8505734, GSM8505735, GSM8505736, GSM8505737, and GSM8505738; and OCM datasets generated in this study for 10 stromal populations). Each dataset was first individually processed in *Seurat*^33^ to remove low quality cells using manual thresholds (nFeature > 1000, nFeature < 7000, nCount > 2000, nCount < 65000, and percent.mt < 5.5), cell doublets using *scDblFinder*^34^ by batch, and finally outlier cells using automatic thresholds (more than 3 MADs away from the median for log(nCount) and log(nFeature), and more than 3 MADs higher than the median for percent.mt). For datasets profiling the stromal compartment, we additionally performed ambient RNA contamination estimation and removal using *SoupX*^35^ prior to doublet detection.

Reference R1 for classifying hematopoietic vs. non-hematopoietic cells was constructed by merging all the above datasets and running *Seurat::NormalizeData*, *Seurat::FindVariableFeatures*, *Seurat::ScaleData*, and *Seurat::RunPCA* with default parameters; the binary labels were determined either by the sample source, or in case where the sample involved both hematopoietic and non-hematopoietic cells, by previously published annotations^3^. *NicheScribe* uses *Seurat’s* anchor-based transfer mapping for classification via *Seurat::FindTransferAnchors* and *Seurat::TransferData* based on precomputed lists of features and 30-dimensional PCA embeddings and returns both predicted labels and prediction scores. A secondary stroma-only reference was generated by applying the first level of *NicheScribe* to reference R1 and retaining cells annotated as ‘non-hematopoietic’ by the classifier. For reference R2.1, we further subsetted cells belonging to one of ten FACS-sorted stromal cell populations and used the hashed labels for training. For reference R2.2, we performed unsupervised clustering using *Seurat::FindClusters* with the smart local moving algorithm and identification of cluster marker genes using *Seurat::FindAllMarkers*. We then performed manual cluster annotation of the R2.2 reference using key marker genes from previous studies^3,12^. A reference UMAP embedding computed with *uwot*^36^ is also provided in order to enable consistent projection of datasets onto a standardized embedding space using *Seurat::IntegrateEmbeddings*.

We benchmarked several widely-used classifiers for scRNA-seq data, including *Azimuth*^37^, *scmap-cell*, *scmap-cluster*^38^, *SingleR*^39^, and *Symphony*^40^. We also compared variants of the *Seurat* query mapping pipeline, including CCA and SCT-based integration. To our knowledge, there is no previous method for annotating cells using a reference specific for murine BM stromal cells, so we instead devised a five-fold cross-validation strategy using our own data to assess performance. We combined 6 CM and Endo stroma and 10 OCM hashing scRNA-seq datasets and randomly split the merged dataset into five parts while ensuring each part contained a similar proportion from each source dataset. For each method and each of the five parts, we reprocessed the other four parts from scratch, used them to trained a new classifier, and predicted cell type labels for the left-out part. Using the hashing labels as ground truth, we computed the accuracy, precision, recall, and F1 scores of these classifiers. The *Seurat*-based implementation of *NicheScribe* was designed based on the results of these tests. *NicheScribe* is available as a R package from github.com/RabadanLab/NicheScribe.

### iMSC-L score

To derive an iMSC-L score, we analyzed two OCM-processed scRNA-seq datasets of pIC-exposed CM MSC-L and Endo MSC-L cells, processed similarly as described above. Together with the two scRNA-seq datasets of normal CM MSC-L and Endo MSC-L, we used *Seurat::FindMarkers* to find differentially expressed gene features between normal MSC-L and iMSC-L. The top 100 genes differentially upregulated in iMSC-L based on adjusted *P* value were used as a signature to score *NicheScribe*-annotated MSC-L cells using *UCell::AddModuleScore_UCell*^29^.

### Analysis of previously published datasets

Data from a previous study of BM stromal cells^6^ were downloaded from the Gene Expression Omnibus (GEO) accession GSE128423 (samples GSM3674224, GSM3674225, GSM3674226, GSM3674227, GSM3674228, and GSM3674229) and reprocessed as described above. The original annotation was recreated using lists of marker genes for each cluster, which were applied using *SCINA*^41^. For analysis of spatial transcriptomics of the murine femur^16^, we downloaded data from GSE297119 (sample GSM8984115). The fastq files were obtained and aligned to the Visium Mouse Transcriptome Probe Set v2.0 mm10 using Space Ranger Count v4.0.1. Visium HD spots were binned to 24 μm diameter for further analysis. For deconvolution analysis, we used *CARD*^17^, supplying the R2.2 reference as cell type metadata and with settings minCountGene = 100 and minCountSpot = 5. For comparison of NicheScribe reference data to human stromal cell types, an analyzed Seurat object of human stromal cells^4^ was downloaded directly (GSE253355), and normalized pseudobulk profiles were obtained using the *AggegrateData* function in *Seurat*. Gene names in the murine reference data were converted to human orthologs using the *getBM* function from the R package *biomaRt*^42^ before correlation in gene expression between cell types was evaluated with the function *clustify* from the R package *clustifyr*^43^, using the top 3000 variable genes from the murine reference object. Output from *clustifyr* was scaled between 0 and 1 across NicheScribe cell populations.

### Analysis of stromal perturbation responses

For analysis of stromal perturbations, datasets were downloaded from GEO (5-fluorouracil [5FU]^7^: GSE108892; pIC^3^: GSE276784; *Mpl^W515L^* MPN^18^: GSE228995; AML^6^: GSE128423) and processed as described above. After applying NicheScribe annotations to each dataset, differential abundance of cells in perturbation conditions vs. matched control samples was assessed using *MELD*^21^, using the same approach as described previously^44^. To evaluate MSC-L responses to perturbation, nonhematopoietic cells were subset for each condition and matched control, normalized using *Seurat::SCTransform*, then integrated using *Seurat::SelectIntegrationFeatures*, *Seurat::PrepSCTIntegration*, *Seurat::FindIntegrationAnchors*, and *Seurat::IntegrateData*. After running *Seurat::PrepSCTFindMarkers*, CM MSC-L defined by annotation R2.2 were subset from the main object, and this matrix was used as input for consensus non-negative matrix factorization (cNMF) using the *cNMF* package^25^.

### Quantification and statistical analysis

Data are represented as means ± standard deviations (S.D.), or as violin plots with the center line representing the median, using R studio for molecular data or GraphPad Prism (version 10.4.2) for all other data. Circles on bar graphs represent biological replicates. For experimental data, Student’s t-test was used when 2 groups were compared. Either one-way ANOVA with Tukey’s post-hoc test or Kruskal-Wallis test with Dunn’s post-hoc test was used to compare 3 or more groups. Data collection and analysis were not performed blind to the conditions of the experiments. Approximate sample size was predetermined for most experiments based on our experience with a range of mouse inflammatory and regenerative perturbations^9^, where a standard deviation of less than 10% can be expected for most measurements. To show a difference of at least 10%, with α □= □ 0.05 and 80% power, 6–11 independent biological replicates are required per group.

## SUPPLEMENTAL INFORMATION TITLES AND LEGENDS

**Document S1.** Figures S1-S7 with figure legends.

**Table S1.** Summary of cNMF factors identified in MSC-L across perturbations.

**Table S2.** Gene loading scores for cNMF model in MSC-L across perturbations.

**Table S3**. Genes used in iMSC-L score.

