## Supplementary material for "NicheScribe: consistent and generalizable annotation of stromal cells in the murine bone marrow niche": Table 1

**Table 1. Correspondence of stromal cell annotations across NicheScribe layers and studies.**

|  | **This study** | | **Mouse** | | | **Human** |
| --- | --- | --- | --- | --- | --- | --- |
| **Location** | **NicheScribe** transcriptomic annotation (R2.2) | **NicheScribe**  FACS prediction (R2.1) | Baryawno *et al*., 2019  (ref. 6) | Kusumbe *et al*., 2014 (ref. 12) | Mohanakrishnan *et al*., 2024  (ref. 13) | Bandyopadhyay et al., 2024  (ref. 4) |
| Central marrow (CM) | Leptin receptor (LepR)-expressing mesenchymal stromal cell (MSC-L) | CM MSC-L | LepR MSC | – | – | APOD+ MSC  THY1+ MSC  Adipo-MSC |
|  | Type L sinusoidal EC (lsEC)1 | CM sinusoidal EC (SEC) | SEC | Type L EC | Bone marrow EC (bmEC) | SEC |
| Mixed (Mix) | Osteoblast-biased (Osteo) MSC-L | CM MSC-L | LepR MSC/  Osteolineage cell (OLC)1 | – | – | APOD+ MSC  Osteo-MSC |
|  | Type H EC (hEC)1 | CM arterial EC (AEC) | SEC/AEC | Type H EC | Metaphyseal EC (mpEC) | SEC |
| Endosteal (Endo) | Adipocyte-biased (Adipo) MSC-L | Endo MSC-L | LepR MSC | – | – | APOD+ MSC  THY1+ MSC |
|  | Sca-1-expressing MSC (MSC-S) | Endo MSC-S | Fibroblast 1 | – | – | Fibroblast-biased  (Fibro-)MSC |
|  | Fibroblast progenitor (FPr) | Endo MSC-S | Fibroblast 2 | – | – | Fibro-MSC |
|  | Osteoblast progenitor (OPr)1 | Endo multipotent mesenchymal progenitor (mMPr) | Fibroblast 3  Fibroblast 5 | – | – | Fibro-MSC |
|  | OPr2 | Endo mMPr/CD24^-^ mesenchymal progenitor (MPr) | Fibroblast 3  OLC2 | – | – | Fibro-MSC |
|  | OPr3 | Endo CD24^-^ MPr | OLC1 | – | – | Osteoblast |
|  | Chondroblast progenitor (CPr)1 | Endo CD24^+^ MPr | Fibroblast 4  Chondrocyte progenitor | – | – | Fibro-MSC |
|  | CPr2 | Endo CD24^-^ MPr/CD24^+^ MPr | Chondrocyte | – | – | Fibro-MSC |
|  | lsEC2 | Endo SEC | SEC | – | – | SEC |
|  | hEC2 | Endo SEC/AEC | AEC | – | – | SEC |
|  | Arterial EC (aEC) | Endo AEC | AEC | Arteries | aEC | AEC |
|  | Type R EC (rEC) | Endo AEC | AEC | – | rEC | SEC |
|  | Pericyte | Endo CD24^-^ MPr | Pericyte | – | – | Vascular smooth muscle cell (VSMC) |
