## Supplementary Figures 1-6 for "NicheScribe: consistent and generalizable annotation of stromal cells in the murine bone marrow niche"

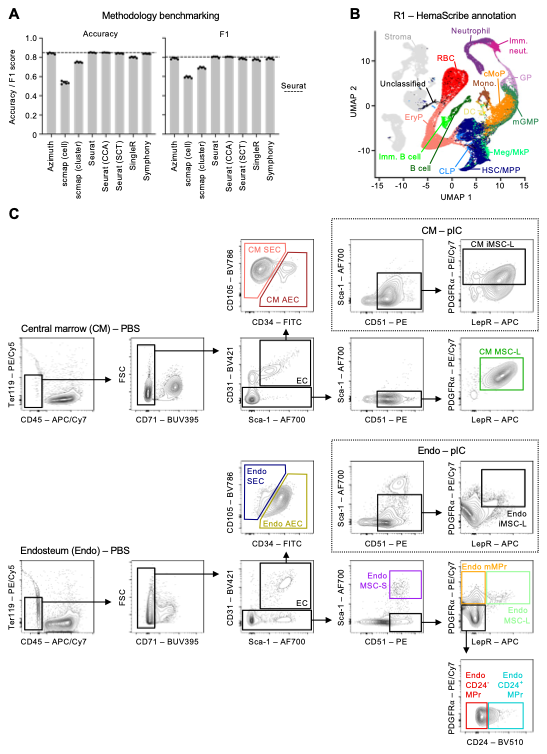

**Figure S1. *Seurat* outperforms other methods for niche cell annotation, related to Figure 1.**

(A) Bar charts showing comparison of indicated methods for annotation of stromal cells. Points represent parts downsampled from reference dataset with mean ± S.D.

(B) UMAP showing reference data for R1 annotation layer, with hematopoietic cells colored by output from HemaScribe: CLP, common lymphoid progenitor; cMoP, common monocyte progenitor; DC, dendritic cell; EryP, erythroid progenitor; GP, granulocyte progenitor; HSC/MPP, hematopoietic stem cell/multipotent progenitor; Imm. B cell, immature B cell; Imm. Neut., immature neutrophil; Meg/MkP, megakaryocyte/megakaryocyte progenitor; mGMP, multipotent granulocyte macrophage progenitor; Mono., monocyte; RBC, red blood cell.

(C) Representative flow cytometric plots showing gating schemes for indicated stromal cell populations used for R2.1 on-chip multiplexing reference: CM, central marrow; EC, endothelial cell with AEC, arterial EC, and SEC, sinusoidal EC; Endo: endosteal; (m)MPr, (multipotent) mesenchymal progenitor; MSC-L, Leptin receptor-expressing mesenchymal stromal cell with iMSC-L, inflammatory MSC-L; MSC-S, Sca-1-expressing mesenchymal stromal cell. Dotted line insets show the gating scheme for iMSC-L isolation in polyinosinic:polycytidylic acid (pIC)-treated mice.

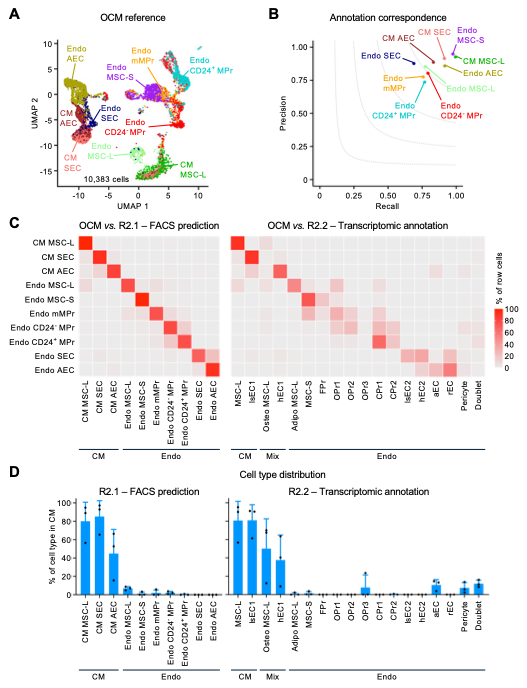

**Figure S2. NicheScribe R2.1 annotation has high accuracy compared to ground truth, related to Figure 1.**

(A) UMAP showing cells isolated by FACS for on-chip multiplexing (OCM), colored by sorted population.

(B) Scatter plot showing precision and recall for indicated populations, comparing NicheScribe R2.1 annotation with ground truth OCM cell type identity.

(C) Heatmaps showing correspondence between OCM reference ground truth (rows) and either NicheScribe R2.1 annotation (left) or R2.2 annotation (right). Rows are scaled to 100%. Adipo, adipocyte-biased; aEC, arteriolar endothelial cells; CPr, chondrocyte progenitor; FPr, fibroblast progenitor; hEC, type H endothelial cell; lsEC, type L sinusoidal endothelial cell; OPr, osteoblast progenitor; Osteo, osteoblast-biased; rEC, type R endothelial cell.

(D) Bar charts showing proportion of indicated cell types found in central marrow (CM), endosteal (Endo) or both locations (Mix). Points represent biological replicates, each composed of 2-4 mice per group, with paired CM and Endo scRNA-seq samples. Bars show mean ± S.D.

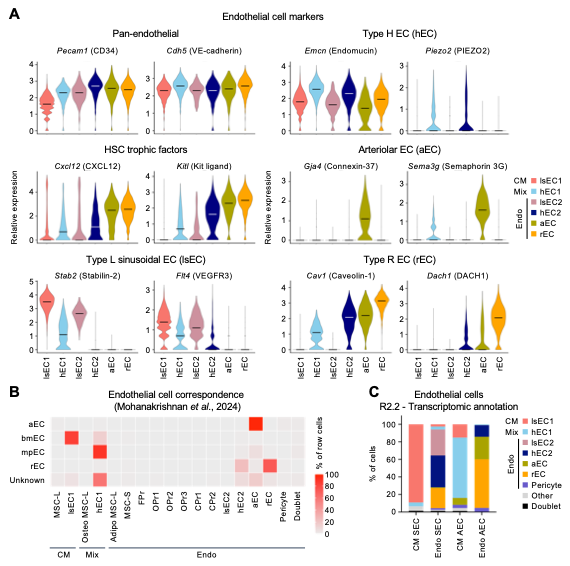

**Figure S3. NicheScribe R2.2 annotation incorporates endothelial cell heterogeneity, related to Figure 1.**

(A) Violin plots showing expression of indicated genes in different endothelial cell subsets defined in the NicheScribe R2.2 annotation. aEC, arterial endothelial cell; hEC, type H endothelial cell; HSC, hematopoietic stem cell; lsEC, type L sinusoidal endothelial cell; rEC, type R endothelial cell.

(B) Heatmap showing correspondence between endothelial cell types identified in published study versus NicheScribe R2.2 transcriptomic annotation organized by central marrow (CM), endosteal (Endo) or both locations (Mix). Rows scaled to 100%. bmEM, bone marrow EC; mpEC, metaphyseal EC.

(C) Bar chart showing composition of NicheScribe R2.1 OCM reference endothelial cell types (columns) according to endothelial cell types in NicheScribe R2.2 transcriptomic annotation (colors).

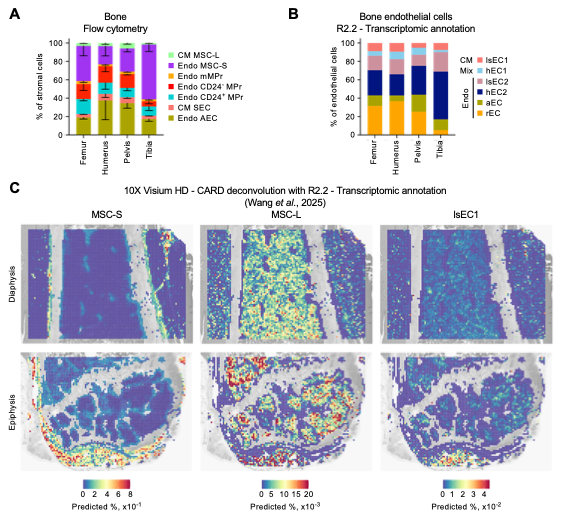

**Figure S4. Spatial localization of stromal cells, related to Figure 2.**

(A) Bar chart showing composition of endosteal stromal cells isolated from indicated bones by flow cytometry. Stacked bars show mean ± S.D. for n=3 mice.

(B) Bar chart showing frequencies of indicated endothelial cell subsets from NicheScribe R2.2 transcriptomic annotation in indicated bones.

(C) Spatial feature plots for selected diaphyseal and epiphyseal regions of published murine femur spatial transcriptomics dataset. Color scale shows predicted composition of spots for indicated cell types using *CARD* deconvolution and NicheScribe R2.2 transcriptomic annotation.

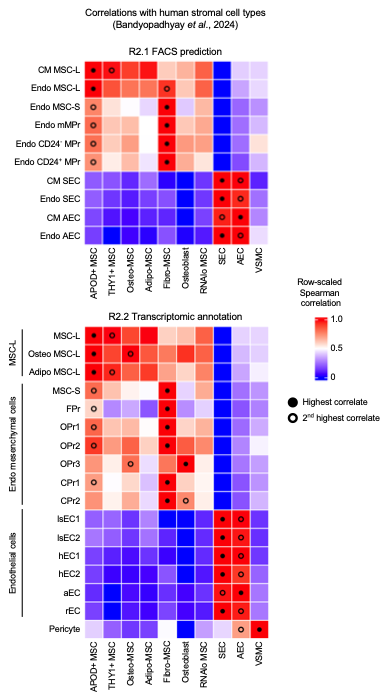

**Figure S5. Conservation of molecular profiles between murine and human stromal cell types, related to Figure 2.**

Heatmaps showing Spearman coefficients for correlations between NicheScribe R2.1 FACS prediction (top) and R2.2 transcriptomic annotation (bottom), scaled between 0 and 1 for each murine population. Black circles indicate most correlated population; open circles show second most correlated population. Fibro, fibroblast-biased; VSMC, vascular smooth muscle cell.

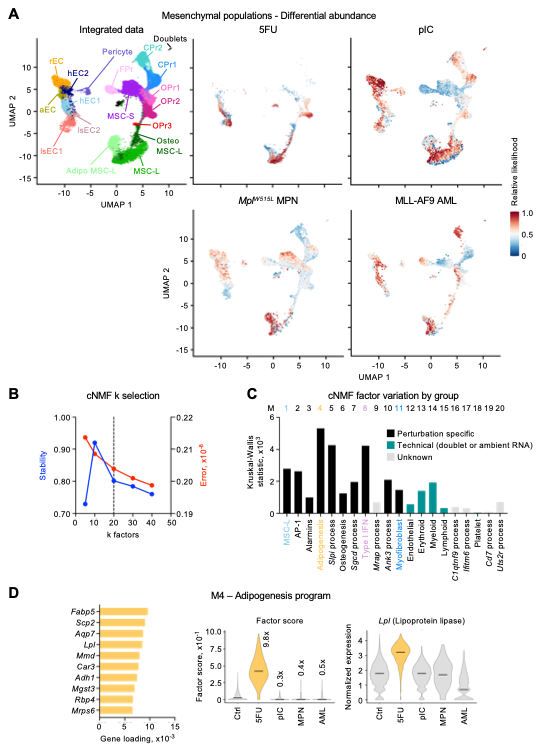

**Figure S6. Changes in stromal cell frequency with perturbation, related to Figure 3.**

(A) UMAP (left) showing cell type composition for all data used in perturbation response comparisons, colored by NicheScribe R2.2 transcriptomic annotation. Feature plots (right) showing relative likelihood of cells by *MELD*, comparing each indicated perturbation to its matched control. 5FU, 5-fluorouracil at day 4; pIC, polyinosinic:polycytidylic acid at day 3; *Mpl^W515F^* MPN, thrombopoietin receptor mutant myeloproliferative neoplasia at 4 weeks; MLL-AF9 AML, MLL-AF9-induced acute myeloid leukemia at 6-10 months.

(B) Line plot showing stability and error for different numbers of k factors in consensus non-negative matrix factorization (cNMF) analysis, with selected k=20 factors named M1-20.

(C) Bar chart showing Kruskal-Wallis (KW) test statistic for comparison of cNMF factor scores across perturbations. Bars are colored according to manual classification.

(D) Description of the M4 Adipogenesis cNMF factor with bar chart showing gene loading scores for top 10 genes (left), violin plots showing factor score annotated with fold change in mean values for each group compared to control (middle), and violin plot showing expression of example gene *Lpl* across groups (right). Lines in violin plots show median values.

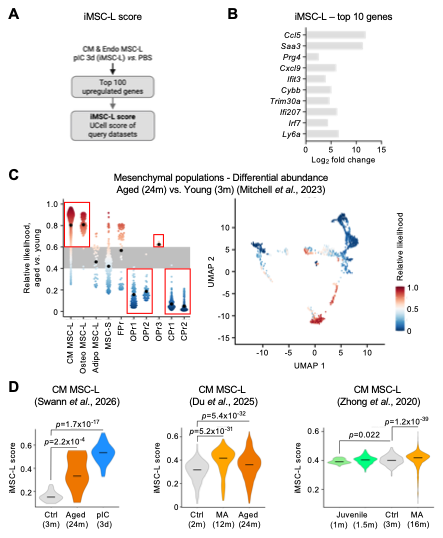

**Figure S7. Emergence of inflammatory MSC-L with aging, related to Figure 3.**

(A) Outline showing generation of a score for inflammatory MSC-L (iMSC-L) activity.

(B) Top 10 genes by adjusted *P* value in the iMSC-L gene signature.

(C) Change in stromal cell frequencies with aging. Beeswarm plot (left) and feature plot (right) showing changes in relative likelihood of indicated mesenchymal cell types or all stromal cells, comparing published scRNA-seq data from aged to young mice. M, month.

(D) iMSC-L score in CM MSC-L from published scRNA-seq data across indicated aging time points. Ctrl, control; d, day; MA, middle-age; pIC, polyinosinic:polycytidylic acid. Lines in violin plots show median values. *P-values* from Kruskal-Wallis test with pairwise Dunn’s test.
